# Low Intensity Multi-Channel Steering TMS Array for Network Level Neuromodulation

**DOI:** 10.64898/2026.08.20.745810

**Authors:** Dexuan Tang, Corey Swenson, Sam Small-Zlochower, Gustav Bizik, Lisbeth Møller Christensen, Thomas Knoesche, Jens Haueisen, Reinhold Ludwig, Guillermo Núñez Ponasso, Gregory Noetscher, Zhi-De Deng, Sergey N. Makaroff

## Abstract

**Objective:** Low-intensity transcranial magnetic stimulation (LI-TMS) is being investigated as a gel-free alternative to transcranial electrical stimulation (tES), but existing systems remain almost exclusively single-channel and cannot electronically steer the induced electric field. We present the design, modeling, and experimental measurement of a wearable whole-head, multichannel, steerable LI-TMS array.

**Methods:** The system comprises a 102-channel conformal coil array with independently controlled drivers capable of arbitrary waveform synthesis, together with a boundary element fast multipole method (BEM-FMM) framework that computes the coil currents required to produce prescribed cortical field patterns. A 12-channel prototype was characterized by coil-current, electric-field, and thermal measurements.

**Results:** The prototype produced a peak primary electric field of approximately 1 V/m measured in air 4 cm from the inner helmet surface. Whole-array modeling attained cortical fields of up to 1.5 V/m, reproduced the field distribution of a clinically validated low-intensity stimulator to within 3%–5%, and demonstrated focal targeting of the dorsolateral prefrontal cortex, simultaneous delivery of electric field to the default mode network nodes, and synthesis of electric fields following the traveling alpha wave.

**Conclusion:** Electronically steerable, whole-head LI-TMS is feasible using accessible microprocessor-controlled power electronics.

**Significance:** The array reaches the cortical field regime of tES without scalp contact or the associated shunting of current through the scalp, offering a route to testing network-level, phaselocked weak-field neuromodulation.

## I. Introduction

Low-intensity (LI) or subthreshold transcranial magnetic stimulation (TMS) refers to stimulation delivered at intensities insufficient to directly elicit neuronal action potentials [1]. Despite operating below the neuronal firing threshold, LI-TMS is being actively investigated for its neurobiological effects in both healthy and pathological conditions, including depression, neural injury and regeneration, abnormal circuit organization, and tinnitus [1]–[7].

Published studies indicate that LI-TMS typically generates electric fields in the brain between roughly 0.01 and a few volts per meter (V/m). Reported values reach 2.2–6.5 V/m within a few millimeters of the coil in rodent preparations [4]–[6], whereas in humans a commercial system induces approximately 0.01–2 V/m [7], depending on hardware design. Collectively, these studies place LI-TMS one to two orders of magnitude below the threshold for directly evoking neuronal action potentials.

Several other low-field techniques developed under different names and design traditions. Pulsed electromagnetic field stimulation (PEMF) systems, which induce cortical electric fields on the order of 0.1 V/m or less [8]; historically designed to maximize magnetic flux density rather than induced electric field. Several PEMF devices have recently been investigated as treatments for depression, including the Re5 system, a single-channel device comprising seven coils driven in phase [9]– [12]; a modern Japanese single-channel micropower system [13]; a rotating-magnet system [14], [15]; and a two-channel PEMF array comprising 20 coils, with ten coils driven in phase and the remaining ten driven in antiphase [16]. Low-field magnetic stimulation (LFMS), which employs MRI-gradient-derived hardware to induce cortical electric fields of comparable magnitude and has been evaluated in randomized, double-blind trials in bipolar and treatment-resistant depression [17]– [19]. These methods differ in coil design, field magnitude, and application, but all are realized in single-channel or fixed-phase hardware; none can electronically steer and induced field or address multiple targets independently.

LI-TMS operates in a low-field regime that is comparable, in order of magnitude, to transcranial electrical stimulation (tES), which induces cortical electric fields of approximately 0.1–1 V/m [20], [21]. Recent approval of Transcranial direct current stimulation (tDCS) for the treatment of major depressive disorder (MDD) indicates the therapeutic potential of neuromodulation within this low electric-field regime [22]– [25]. Notably, tES is already available in multichannel form: commercial high-definition tES platforms provide arrays of independently driven scalp electrodes [26], [27], and multichannel transcranial alternating current stimulation (tACS) has been used to impose prescribed phase relationships across cortical sites, yielding frequency- and phase-dependent modulation of cortical excitability and behavior [28], [29].

It should be emphasized that the evidence for clinical efficacy of LI-TMS remains scarce. Suprathreshold rTMS for depression is supported by large multicenter randomized sham-controlled trials, consensus recommendations, and regulatory clearance [30]–[32]. By contrast, LI-TMS has not been evaluated in adequately powered randomized sham-controlled trials; the available human evidence consists largely of small open-label or single-arm studies, with the bulk of mechanistic support coming from rodent models [1]–[3]. Where related low-field modalities have been tested under randomized, double-blind conditions, reported antidepressant effects have been modest and inconsistent [10], [11], [17]–[19]. The system described here is a hardware feasibility and modeling study; it has not been evaluated in any clinical trial, and no claim of therapeutic efficacy is made.

The most informative account of what weak fields can do instead comes from tACS, which operates in the same cortical field regime as LI-TMS, approximately 0.1–1 V/m, and where the proposed mechanism is not the initiation of action potentials but the biasing of spike timing within ongoing activity. Intracranial recordings in nonhuman primates show that tACS at these amplitudes entrains single-neuron firing to the applied waveform [33], and recordings in awake animals show preferential phase synchronization of cortical neurons to an applied alpha-band waveform [34]. On this account, weak fields act by coupling into, and phase-locking with, the brain’s own oscillations rather than by imposing activity upon them [35], and dose-response estimates place the minimum effective field in the same range that the present array attains [36]. Complementary evidence from rodent LI-TMS preparations points to subthreshold membrane polarization and intracellular calcium signaling, with downstream effects on gene expression and functional connectivity [1], [5], [6].

This account motivates the present hardware and identifies a specific advantage over tACS. If the operative mechanism is phase-locked coupling to endogenous rhythms at fields of order 1 V/m, then what matters is the ability to place fields of that magnitude, with controlled phase and frequency, at several targets at once, which is what a multichannel inductive array provides. In transcranial electric stimulation, however, a large fraction of the applied current shunts through the scalp rather than entering the brain, and further current is diverted through the highly conductive cerebrospinal fluid, so that cortical fields of only about 0.8 V/m are reached at 2 mA [21]. Magnetic induction is not subject to this shunting: the magnetic field traverses scalp and skull essentially unattenuated and induces the electric field directly in tissue. An inductive array can therefore deliver comparable cortical field magnitudes without the scalp losses that constrain tES and, by exploiting constructive interference among many coils, can bias the field toward deeper structures instead of being confined to superficial cortex. Whether weak-field coupling accumulates into durable, clinically meaningful responses in humans remains an open question, and it is the question that steerable, network-level hardware of the kind described here is intended to make experimentally feasible.

LI-TMS hardware remains almost exclusively single-channel. Most practical LI-TMS systems reported to date (cf. [1], [4]–[7]) are based on relatively simple single-channel architectures employing one or, at most, two custom coils driven by a single function generator and power amplifier. To the best of the authors’ knowledge, no steerable or phased-array LI-TMS hardware currently exists that is capable of selectively stimulating multiple predefined cortical or subcortical targets, either sequentially or simultaneously.

This situation contrasts with suprathreshold TMS, for which several multichannel hardware systems have already been realized, including a multi-channel stimulator driving independently controlled coil elements with separately adjustable parameters [37], a five-channel array of overlapping TMS coils [38], and multichannel arrays based on three-axis coil elements [39]. Achieving a multichannel architecture is considerably more challenging for suprathreshold TMS because of the substantially higher power requirements, making the current absence of multichannel LI-TMS systems particularly noteworthy.

The goal of this study is to demonstrate the feasibility of a wearable whole-head, multichannel, steerable LI-TMS helmet based on accessible microprocessor-controlled power electronics and software-defined control. The proposed system generates cortical electric fields of up to 1.5 V/m, provides acceptable thermal performance, and operates in both pulsed and continuous-wave (CW) modes. A 12-channel prototype is experimentally validated through coil-current synthesis, electric-field steering, and temperature measurements. The capabilities of the full array are then demonstrated through four modeling examples of increasing complexity: (i) upscaled replication of the whole-cortex electric-field distribution of the Re5 PEMF stimulator; (ii) focal stimulation of the dorsolateral prefrontal cortex (DLPFC) with varying focality and field strength; (iii) simultaneous stimulation of multiple regions within the default mode network (DMN) [40]–[42], a principal network-level target and biomarker in depression [42]; and (iv) replication of an endogenous traveling alpha wave. The last example is motivated by evidence that cortical alpha and theta rhythms propagate across the neocortex as traveling waves [43], and by the growing interest in synchronizing stimulation to endogenous cortical rhythms [44]–[46].

## II. Materials and Methods

### A. Stimulator concept

The concept of the proposed stimulator array builds upon extensive theoretical studies of multichannel TMS [47]–[50] and originated from our previous research on MEG source reconstruction [51], [52]. During source-localization studies using the Elekta Neuromag helmet, we observed that its magnetometer coils (102 of the 306 sensors) possess several characteristics desirable for multichannel magnetic stimulation, including a relatively large coil size (approximately 30 × 30 mm), dense conformal coverage of the cortex, and a low-profile assembly geometry (Fig. 1a). By the principle of reciprocity, we hypothesized that similar coils, positioned closer to the scalp (Fig. 1b), could be used not only to record magnetic fields in MEG but also to generate time-varying magnetic fields and the associated electric fields within the cortex. Consequently, cortical spatial resolution comparable to that achieved in MEG may be attainable, potentially approaching 10 mm. Although omitting the gradiometer coils is expected to reduce spatial resolution, particularly for superficial cortical targets, MEG source localization based solely on magnetometer channels has nevertheless been shown to provide reasonably accurate localization [51], [52].

**Fig. 1.**
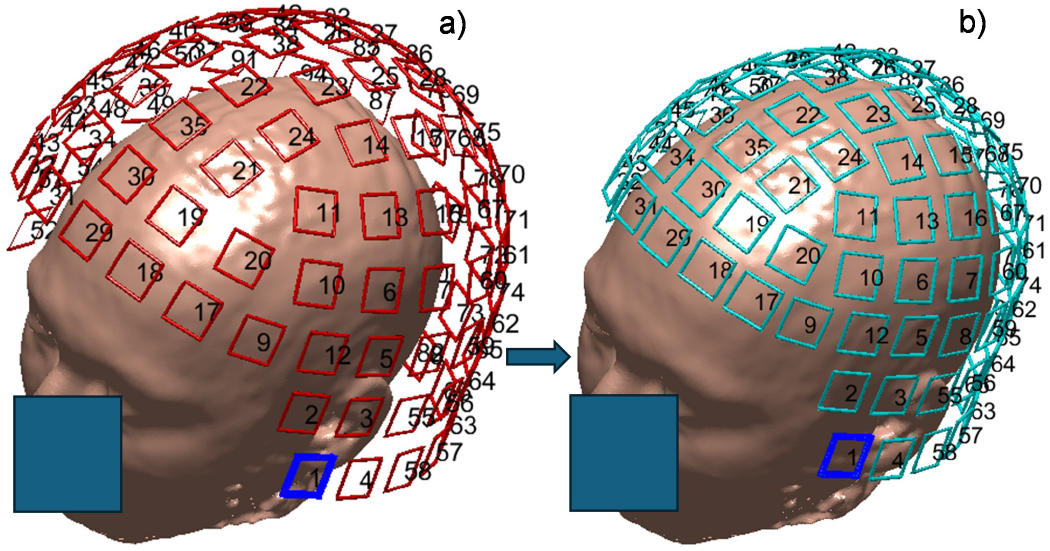
a) Set of Elekta Neuromag MEG magnetometer coils to scale. b) Concept of the present inductive-stimulation array for the same subject to scale.

### B. Stimulator components

The proposed LI-TMS system comprises three principal components (Fig. 2): (i) a rigid, 3D-printed 102-channel coil-array helmet with individually detachable coils and connectors routed along its inner surface to the base; (ii) coil-drive electronics and a battery housed within the red enclosure; and (iii) a host computer that wirelessly controls the current waveform of each coil – including pulse shape, phase, amplitude, and repetition frequency for pulsed operation, or phase and amplitude for sinusoidal operation – to target a specified neural site using a field synthesis algorithm we refer to as beamforming in this paper. Table I summarizes the parameters of the individual coil radiators. The current hardware prototype incorporates 12 fully functional programmable channels, each of which can be connected to any of the 102 array coils or to a selected combination of coils. The remaining 90 channels are at various stages of development.

**Fig. 2.**
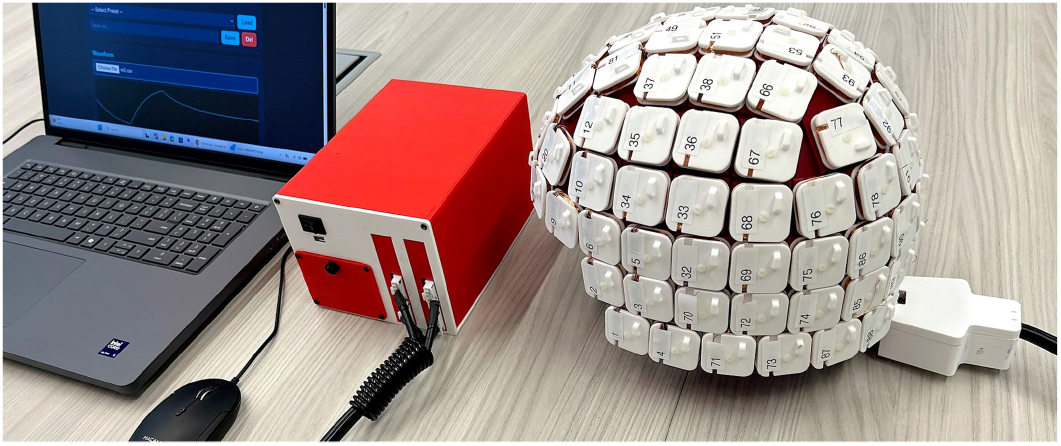
Functioning array prototype used for testing, durability, thermal, and field measurements, as well as experimental and biocompatibility evaluations. The protype is driven by a custom Li-ion battery bank (in the red enclosure)

**TABLE I.** Parameters of an Individual Coil (Current Design, Subject to Change)

| Parameter | Value | Parameter | Value |
| --- | --- | --- | --- |
| Overall dimensions/weight | 33×33×7 mm, ~20 g | DC resistance | ~2.4 $\Omega$ |
| Winding material | 28 AWG enameled copper wire | Inductance at 1 kHz | ~450 μH |
| Number of turns | 104 | Connector | JST-VH, 6 pin, 3.96 mm pin pitch |
| Coil form material | SLS printed PA12 | Power loss | 1.2 W for sine wave with 2 A pk-pk |

### C. Power electronic hardware

The key hardware elements are driver printed circuit boards (PCBs; Fig. 3), each providing six independent, software-controlled coil channels (Fig. 3b). Software-defined control of every coil current enables beamforming across the cortex, limited only by the number and orientation of the coils. Each PCB hosts six waveform generators controlled by a single STM32 microcontroller (mounted on the reverse side of the board) and communicates with other boards via a Controller Area Network (CAN) bus. Arbitrary waveform synthesis is achieved using MOSFET (Nexperia BUK9M34-100E) full H-bridges with current feedback provided by Allegro CT4022-A24BSN8 sensors. The waveform generation scheme is similar to that used in PWM motor controllers [53] and temporal interference or conventional TMS systems [54], [55]. An 80 kHz PWM signal is regulated by a 40 kHz PID control loop, with current feedback continuously adjusting the duty cycle to match the prescribed waveform. Current is sensed and regulated independently in every coil, so that each channel corrects its own duty cycle in closed loop against its measured current rather than against an assumed coil impedance. Mutual inductance between neighboring coils, which in a densely packed array would otherwise perturb the individual currents and distort the synthesized field pattern, is therefore rejected as a disturbance, and the current delivered by each coil tracks its commanded waveform as closely as the hardware permits. We emphasize that 80 kHz is the switching carrier used to synthesize the coil current and is not a stimulation frequency; the delivered waveforms are confined to the sub-kilohertz range, and the carrier itself is suppressed by the coil, as quantified below. Operating at 24 V, the current prototype supports peak coil currents of up to 9 A and a maximum slew rate of 0.05 A/*µ*s (approximately 0.1% of the dI/dt at 100% MSO of conventional suprathreshold TMS). The driver-board specifications are summarized in Table II.

**Fig. 3.**
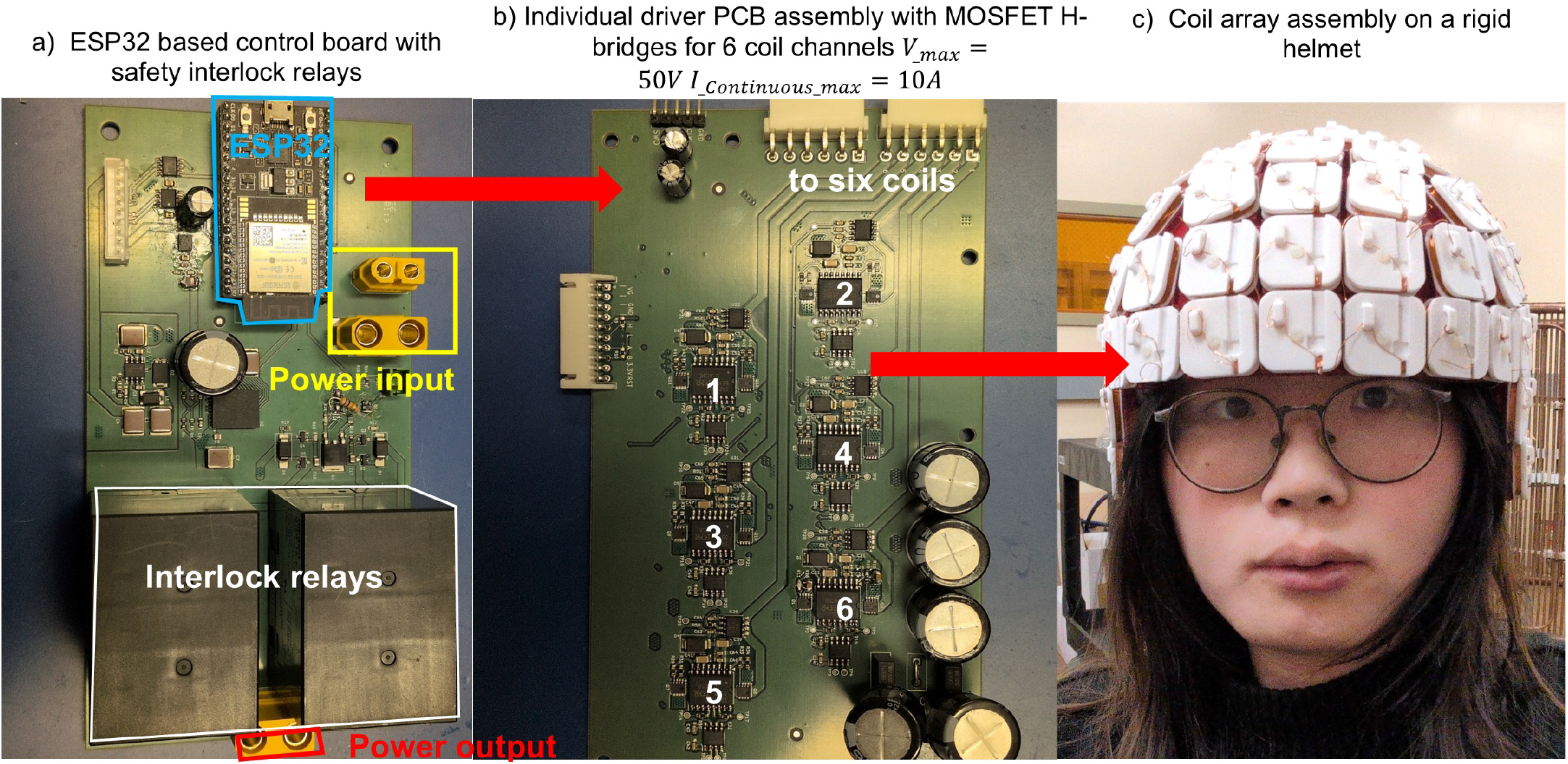
Electronics and communication hardware. a) ESP32 based control board. Interlocking relays only provide power when all systems are functional and safe. b) Driver PCBs with six channels (front side). Each channel is driving one coil of the helmet in c). Subject in c). is one of the authors.

**TABLE II.** Driver Board Specifications.

| Parameter | Value | Parameter | Value |
| --- | --- | --- | --- |
| Overall board dimensions | 165×92×25 mm | Analog current sensor bandwidth | 500 kHz per channel |
| Maximum input voltage | 50 V | ADC resolution | 14 bits per channel |
| Maximum peak current | 10 A per channel | ADC current resolution | 3.6 mA/bit per channel |
| CAN bus speed | 1 Mbit/s | PWM frequency | 80 kHz per channel |
| Analog current sensor resolution | 55 mV/A per channel | Control loop frequency | 40 kHz per channel |

The coil inductance and resistance of Table I are deliberate design decisions With *L* ≈ 450 *µ*H and *R* ≈ 2.4 Ω, the coil time constant is *τ* = *L/R* ≈ 188 *µ*s, approximately fifteen times the 12.5 *µ*s PWM period, so the coil acts as a first-order low-pass filter that removes the carrier from the delivered current. For unipolar switching from the 24 V rail the worst-case peak-to-peak ripple occurs at 50% duty cycle and is Δ*I*_pp_ = *V/*(4*Lf*_PWM_) = 24*/*(4 · 450 *µ*H · 80 kHz) ≈ 0.17 A, below 2% of the 9 A peak current; a proportionally smaller inductance would raise it in inverse proportion. The same inductance also bounds the attainable slew rate at *dI/dt* ≤ *V/L* ≈ 0.053 A/*µ*s, which is what limits the prototype to the 0.05 A/*µ*s quoted above.

### D. Communications hardware and software

Fig. 4a presents a high-level block diagram of the major control hardware and software components. Communication hardware, storage, and execution units are highlighted in gray. Each power driver PCB is controlled by an STM32 microcontroller (mounted on the reverse side of the board; Fig. 4b), which manages six independent channels. All STM32 controllers communicate with a single ESP32S3 (Fig. 4a,b) via a Controller Area Network (CAN) bus. The ESP32S3 provides communication with the host computer either wirelessly through a Wi-Fi hotspot and web interface or via a wired connection. To date, two STM32 controllers and the ESP32S3 have been programmed and validated.

**Fig. 4.**
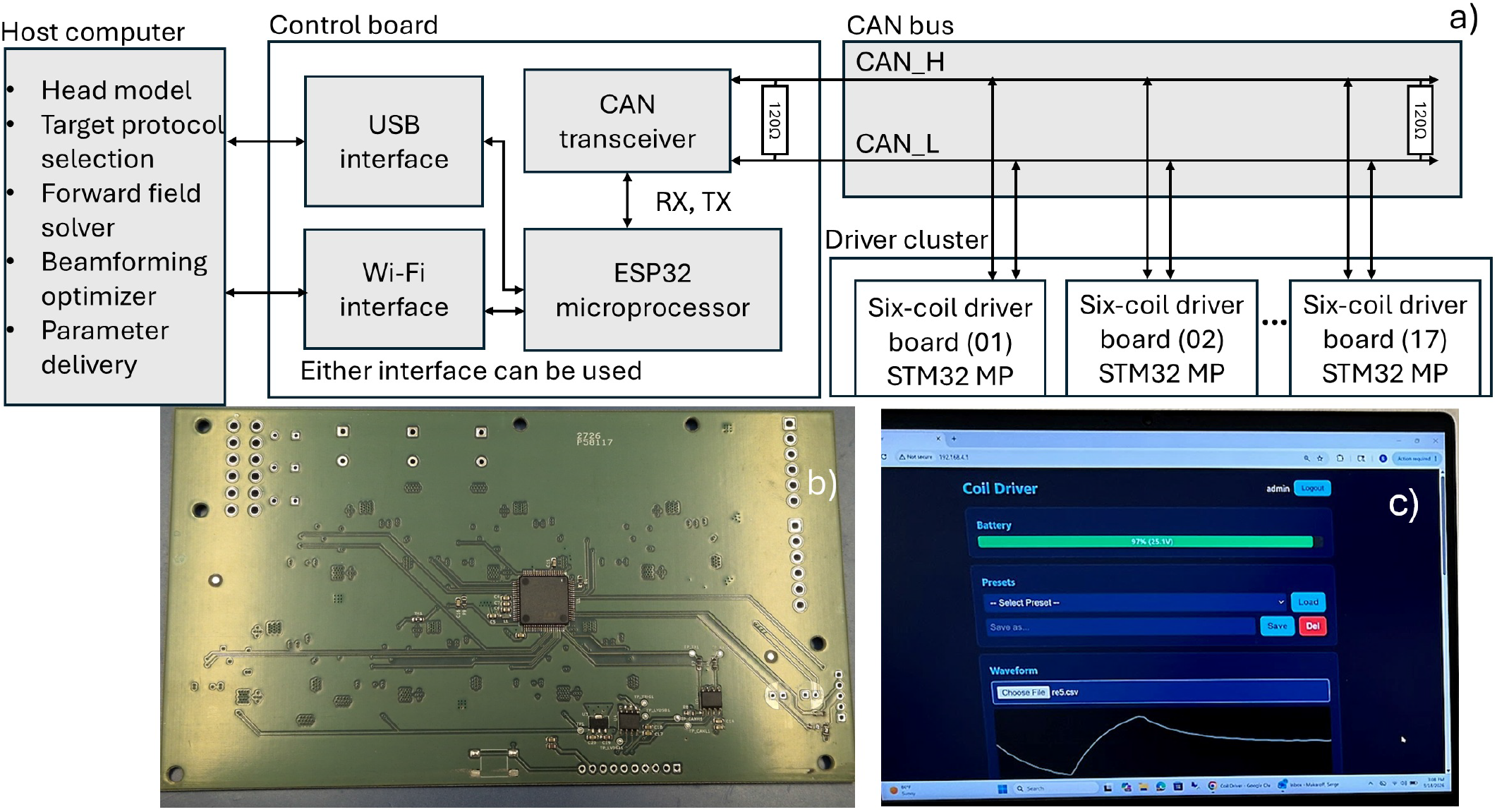
Top-level communications block diagram. The wired USB interface to the host computer is optional; the current design uses a Wi-Fi interface.

An initial version of the control software has also been developed for both the microcontrollers and the host computer. Its primary functions are (i) arbitrary waveform generation for each channel (Fig. 4c) and (ii) beamforming, including the generation of static or dynamic stimulation protocols that prescribe the magnitude and/or polarization of any component of the electric or magnetic field.

### E. Beamforming software

Array beamforming is accomplished on the laptop workstation via two-way communications, as it requires more involved computations. The perquisites for this are (i) helmet-to-head registration transform obtained, for example, using four rigid, fixed fiducial markers on the outside of the helmet and at least the three head fiducials – nasion, LPA, and RPA measured individually and; (ii) an individual head segmentation in *.stl format obtained from a T1-weighted MRI scan of the patient using, for example, the popular *headreco* pipeline [56].

The computational method is the boundary element fast multipole method or BEM-FMM. BEM-FMM [57]–[60] is an iterative method that can serve as a powerful alternative to FEM and classical BEM [58]. While classical BEM approaches use the surface potential as the fundamental unknown, the BEM-FMM uses a charge-based formulation coupled with the fast multipole method (FMM) [61]. As an iterative method, the BEM-FMM does not create a system matrix, so it is able to compute fast solutions using highly complex and realistic meshes such as, for example, the 40-compartment head models of Sim4Life segmentation [62].

Algorithm for spatial beamforming is as follows. The input data is the desired electric field distribution on the head interfaces or in a volume. This could be field magnitude, a normal-to interface electric field, or a full vector electric field. For example, the target field magnitude on a selected Region of Interest (ROI) on the white matter interface (just outside) can be chosen as 0.5 V/m while it is set to zero anywhere else. The output is given by the desired coil currents. BEM-FMM is used to precompute the forward solution of the coil array. An *m*-th coil is energized by setting its current *m* to 1 A and the currents of all other coils to 0. The resulting electric fields on brain interfaces (**E**, 3 × *N*_*faces*_) are computed and stored. These fields form 102 global vector basis functions for the electric field. Once the basis functions are computed, we use the target information by specifying the desired field strength in target interface regions, (**E**_*target*_, × 3 *N*_*faces*_) based on the input data above. Finally, the target (for example, vector) field **E**_*target*_ on brain interfaces is expanded into these 102 basis functions with 102 unknown scalar coefficients – the desired coil currents *m*. After forming a dot product with every basis function, this representation leads to a system of 102 linear equations for 102 unknown coil currents *m*. Such a method is closely related to the Moore–Penrose pseudoinverse or minimum-norm estimation, similar to methods employed in EEG/MEG source reconstruction [63], [64]; it is also similar to the method first used in Ref. [47] and in [65]. A constrained optimization subject to limits on dI/dt is performed.

### F. Measurement setups

#### Measuring electric fields in air

Twelve frontal array channels (17, 18, 29, 31, and 33 in the first row; 18, 20, 19, 23, 22, 26, and 25 in the second row) were driven with the same current waveform, differing only in amplitude and phase. The objective was to achieve focal stimulation of a region of interest (ROI) within the left dorsolateral prefrontal cortex (DLPFC) and compare the measured and simulated electric fields. The induced electric field was measured using a calibrated toroidal electric-field probe constructed as described in [66] and shown in Fig. 5a. This Rogowski-type sensor employs a toroidal winding that cancels the magnetic flux threading the ring, making the probe output proportional to the second time derivative of the normal electric field passing through the toroid. For a fixed current waveform, the output voltage is therefore directly proportional to the local peak-to-peak electric-field magnitude, with the same proportionality constant at all measurement locations. The probe output was filtered with a first-order analog low-pass filter (1.5 kHz cutoff) to suppress the 80 kHz PWM switching ripple and prevent aliasing during digitization, followed by a sixth-order Butterworth low-pass filter (40 kHz cutoff) implemented in software. Signals were acquired with an Agilent DSO6034A oscilloscope, and the peak-to-peak probe voltage was averaged over 15 periods at each measurement point. Electric-field vectors were mapped in air beneath the 12-coil prefrontal montage by orienting the directional probe successively along the three Cartesian axes and translating it on a 1 cm rectangular grid comprising 783 observation points (Fig. 1e) using a three-axis positioning stage (Fig. 5a). The measured electric-field magnitudes were then compared with the corresponding simulations.

**Fig. 5.**
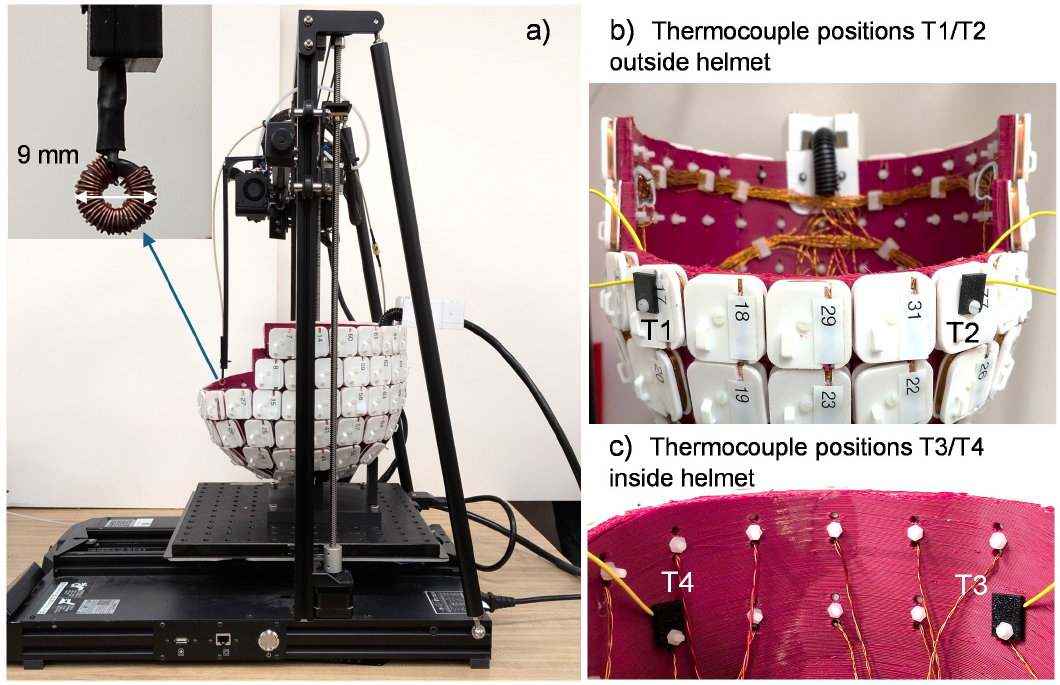
Measurement setups for a) vector electric field in air and; b) for temperature measurements inside and the helmet. Twelve most frontal array channels (17, 18, 29, 31, 33 in the first row and 21, 20, 19, 23, 22, 26, 25 in the second row) were driven with the same current waveform and differed only in amplitude/phase.

#### Measuring array heating

The stimulation protocol described above was used for temperature measurements at different electric-field strengths using a four-channel Extech SDL200 data-logging thermometer. As shown in Fig. 5b, two thermocouples were attached to the left (T1) and right (T2) sides of the coil to monitor peak coil heating. The remaining two thermocouples were attached to the inner helmet surface (Fig. 5c): one above the left DLPFC (T3) to monitor the maximum temperature rise inside the helmet, and the other above the right DLPFC (T4) to monitor the overall helmet temperature. All measurements started from a baseline temperature of 25^*◦*^C. Temperature rise was recorded for both pulsed and continuous-wave (CW) stimulation at identical peak-to-peak electric-field levels.

## III. Results

### A. Measurement results

#### Generating arbitrary waveforms by a single-channel coil driver

As two examples of arbitrary waveform synthesis, we first replicated the current waveform of the Danish Re5 PEMF neurostimulation device [8], [9] using a single array channel at a substantially higher power level. Fig. 6a compares the original Re5 waveform (blue) with the synthesized waveform (red). The slight positive/negative asymmetry was reproduced with a normalized 2-norm error below 5%, while increasing the peak-to-peak current amplitude by a factor of 17. As a second example, we synthesized a widely used theta-burst waveform [67], increasing the peak dI/dt from 0.017 to 0.05 A/*µ*s. Fig. 6b compares the target and measured waveforms, demonstrating accurate reproduction at the higher slew rate. The positive phase of the theta-burst pulse is 1 ms, compared with 2.25 ms for the Re5 waveform.

**Fig. 6.**
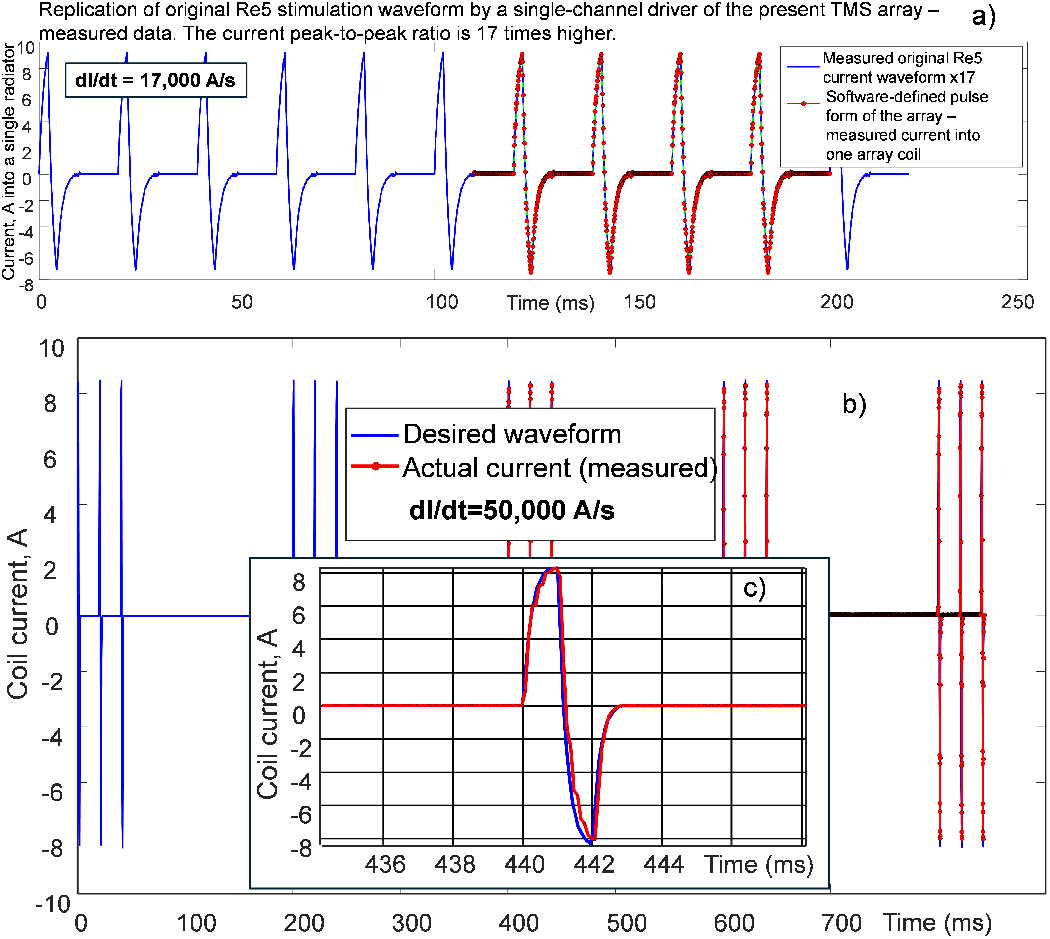
a) Blue – measured current waveform delivered by the authentic Re5 device from their power unit, additionally multiplied by the factor of 17. Red – software-defined and then measured current waveform delivered by one channel of the present driver of the array. b,c) Blue–target(desired) software-defined current waveform of the theta-burst stimulation with 5 Hz repetition frequency. Red – actual, measured current waveform. Inset c) shows the zoomed in individual pulse.

#### Comparison between measurements and computations for targeting left DLPFC

To demonstrate the targeting capability of the 12 measured array channels, the experimental setup described in Section 2.6 was used. The headreco segmentation of Human Connectome Project [68] subject 110411 was registered to the CAD helmet model (Fig. 7a), and the beamforming method of Section 2.5 was applied to target a 3-cm-wide region of interest (ROI) within the left dorsolateral prefrontal cortex (DLPFC) on the white-matter interface (all head compartments were replaced with air in this experiment). The resulting coil-current distribution is shown in Fig. 7a, and the corresponding electric-field magnitude on the white-matter interface is presented in Fig. 7b.

**Fig. 7.**
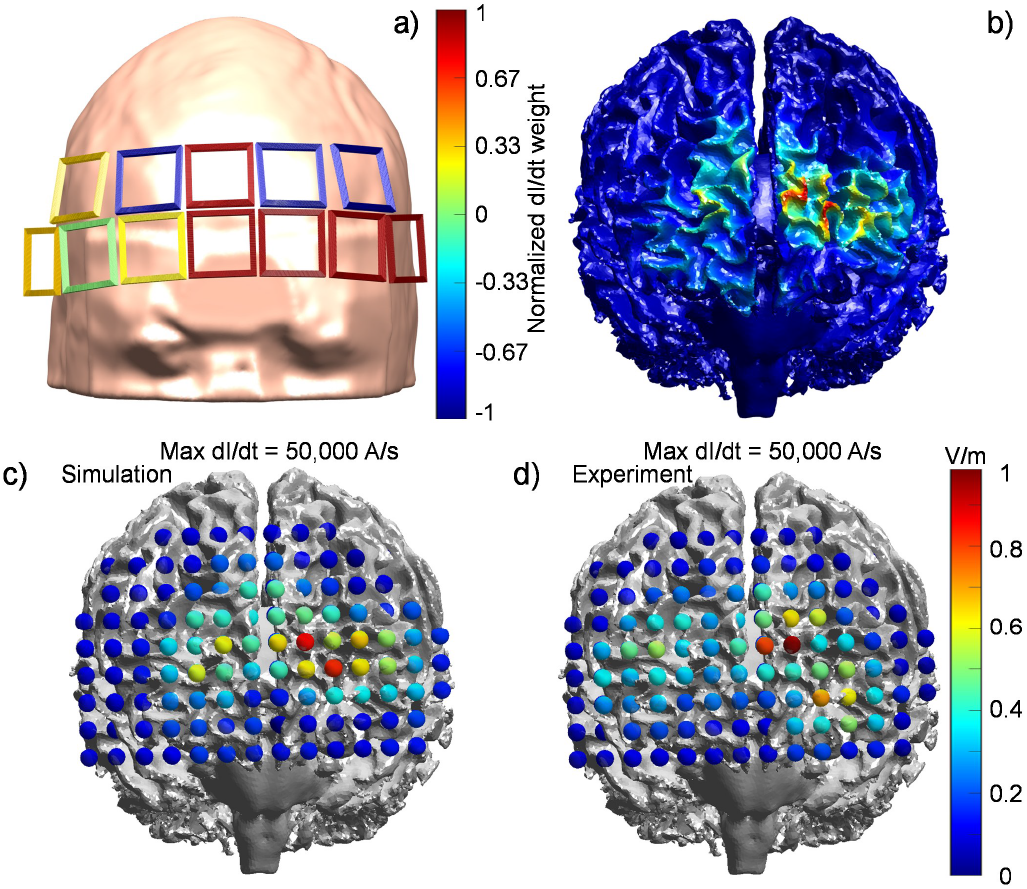
a) Connectome subject 110411 registered within the helmet; only coils used experimentally are shown in this panel to scale. Different colors indicate normalized coil current weights obtained as a result of targeting left DLPFC performed numerically. b) Magnitude of electric field just above (outside) the white matter interface predicted by the beamforming method of Section 2.5. c,d) Simulation and measurement results, respectively, for the grid points located in the vicinity of the white-matter interface.

The synthesized current distribution was implemented experimentally using the pulse waveform of Fig. 6c with a peak dI/dt of 0.05 A/*µ*s. Figs. 7c,d compare the simulated and measured electric-field magnitudes near the white-matter interface. Only the observation points closest to the interface are shown. The agreement between experiment and simulation is within 25% (2-norm). At a distance of 4 cm from the inner helmet surface (the auxiliary white-matter surface), the measured peak electric-field magnitude slightly exceeded 1 V/m (Fig. 7d). At 1 cm from the inner helmet surface, the peak field reached 1.83 V/m in simulation and 1.75 V/m experimentally.

#### Coil heating measurements

To assess the extent of coil heating, we employed the scaled Re5-NTS pulsed stimulation protocol shown in Fig. 6a, which features a relatively long current pulse, as well as a continuous-wave (CW) sinusoidal waveform at 1 kHz, both operating during 10 min. Fig. 8 presents the temperature measurements obtained at the four probe locations shown in Fig. 5b,c for different peak-to-peak coil-current values. No any form of air cooling was employed. The upper-left panel corresponds to the original Re5 current amplitude.

**Fig. 8.**
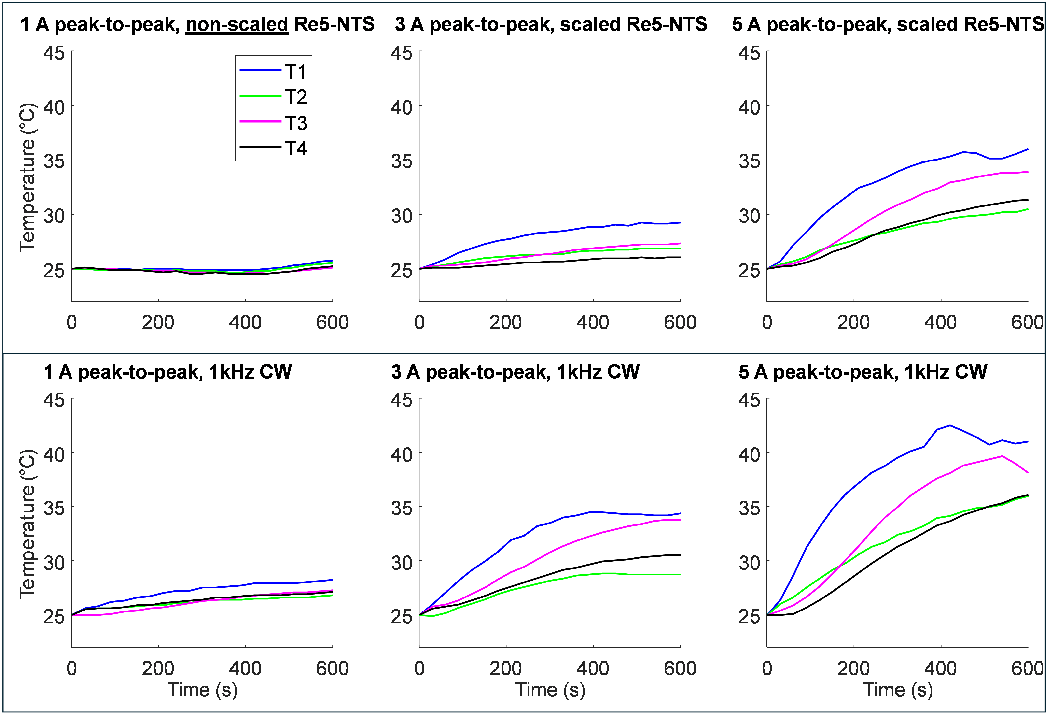
Extent of coil heating at different current strengths and wave-forms.

### B. Modeling results

These results are intended to demonstrate the capabilities of the proposed array for a variety of beamforming tasks. In the examples that follow, all 102 array channels were assumed to be active (instead of 12); whole-head beamforming was performed using realistic head models and realistic tissue-conductivity values.

#### Example #1. Replication of electric and magnetic fields for existing low-power magnetic stimulator - the Danish Re5 system [8], [9]

Fig. 9a,c shows electric and magnetic fields, respectively, of the Re5 stimulator on a white matter interface of a subject from Ref. [8] augmented with SimNIBS conductivity properties. A full vector-field optimization with the above minimum-norm method was used to closely match these fields with the present multi-coil array. Fig. 9b,d shows the corresponding synthesized results for the electric and magnetic field, respectively. The 2-norm difference between the two fields distributions for the white-matter interface does not exceed 3-5% in both cases(!). It should be noted that the E- and B-fields were optimized by using two independent solutions, not simultaneously. It should also be noted that the Re5 cortical E-fields appear to be very small (≤ 0.05 V/m) for the bulk of the brain.

**Fig. 9.**
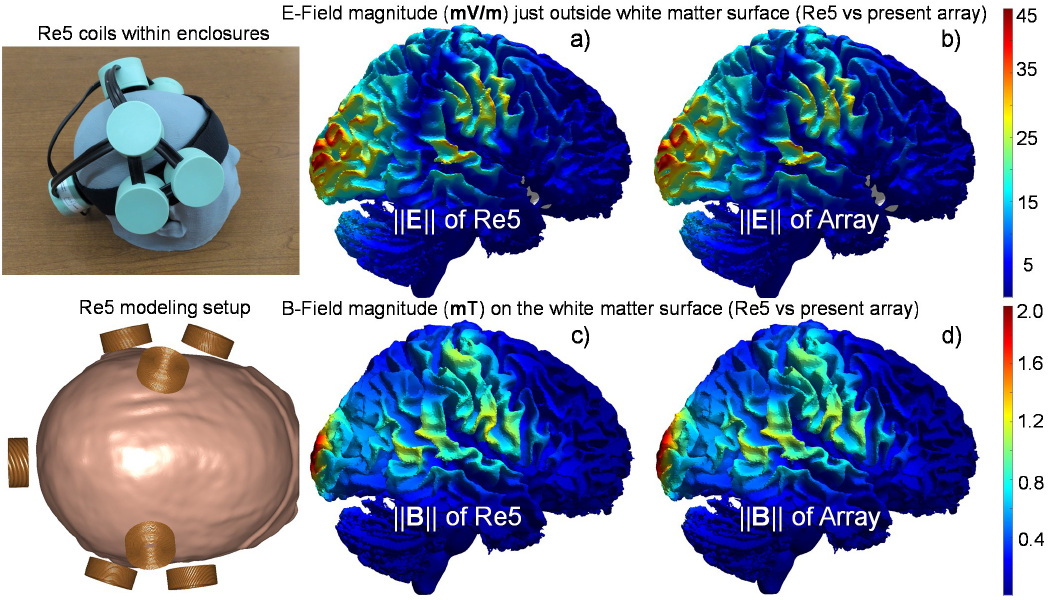
a), c) Authentic electric and magnetic fields of the Re5 stimulator, respectively. b), d) Synthesis of the same fields with the present array.

#### Example #2. Stimulation of a selected cortical region (DLPFC) with varying degrees of focality and field strength

Fig. 10a shows a typical volumetric electric-field distribution obtained by beamforming on a logarithmic scale. The peak cortical field approaches 0.35 V/m when the individual-coil dI/dt is limited to 0.017 A/*µ*s and approximately 1 V/m for the experimental limit of 0.05 A/*µ*s (Fig. 7). At the higher slew rate, however, the current hardware limits the positive pulse duration to approximately 1 ms (cf. Fig. 6c). Figs. 11a–c illustrate targeted stimulation of the left dorsolateral prefrontal cortex (DLPFC) – the principal TMS target for depression treatment [22], [69] – under different beamforming constraints, while Figs. 11d–f show the corresponding normalized coil-current distributions.

**Fig. 10.**
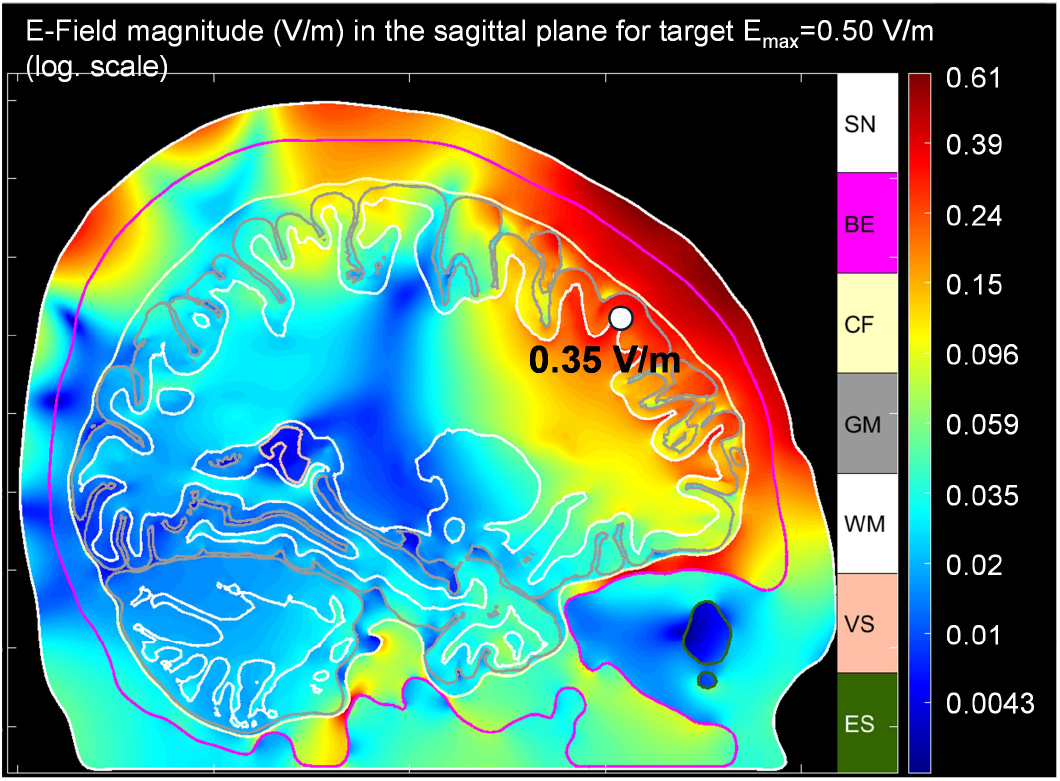
Volumetric electric field-magnitude distribution corresponding to whole-array left DLPFC targeting and the protocol of Fig. 6a (dI/td =0.017 A/***µ***s). Note the logarithmic scale.

**Fig. 11.**
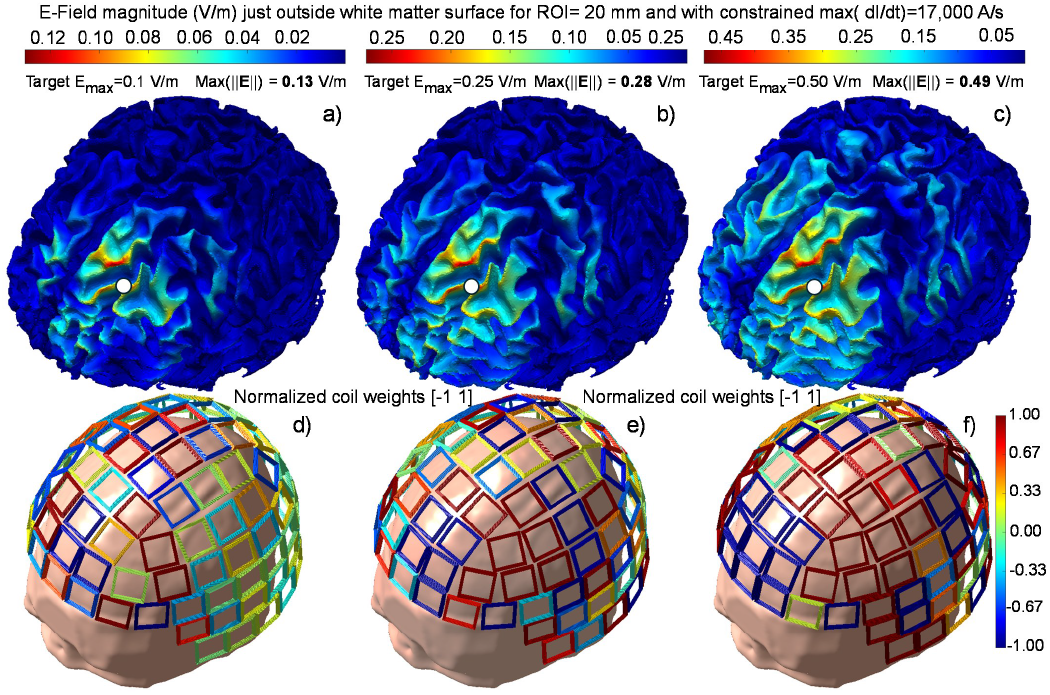
a,b,c) Electric field magnitude distributions just outside the white matter surface for left DLPFC targeting at different target field strengths over a 2 cm wide ROI. d,e,f) Normalized coil current distributions in every case.

The results were obtained using constrained beamforming with the individual-coil dI/dt limited to 0.017 A/*µ*s, still 17 times greater than that of the original Re5 PEMF pulse. The 2-cm-diameter ROI is centered at the white circle in Figs. 11a–c. The highest focality is achieved for the lowest target field (0.1 V/m, Fig. 11a), while increasing the target field reduces focality, although it remains satisfactory, consistent with previous suprathreshold TMS modeling [65]. Interestingly, the optimized coil-current distribution evolves from an apparently random pattern dominated by destructive interference (Fig. 11d) to a more structured configuration (Fig. 11f), where the array effectively forms a large figure-8 coil with two oppositely driven windings.

#### Example #3. Simultaneous stimulation of the default mode network (DMN)

The DMN ([40]–[42]) is considered one of the principal network-level targets and biomarkers in depression research and treatment, including TMS [42]. Fig. 12a shows an fMRI meta-analysis map of the DMN derived from 777 neuroimaging studies in a near-midsagittal plane [[70], [71]. After approximate registration by scaling and aligning the brain models, the corresponding downloadable 3D fMRI voxel mask [71] was used as the beamforming target for the head model shown in Fig. 12b, where regions with highlighted voxel will be optimized for maximum E-field amplitude, while other region’s E-field amplitude will be minimized, within the limitation of the solver and hardware. Constrained optimization with a maximum individual-coil dI/dt of 0.017 A/*µ*s was performed using the Sim4Life head segmentation [62] and its default conductivity values. Figs. 12b,c show the resulting electric-field distributions, displayed on logarithmic and linear scales, respectively. Although the computed field pattern is physically plausible and compatible with network-level modulation [71], further improvements in cortical targeting efficiency are still required.

**Fig. 12.**
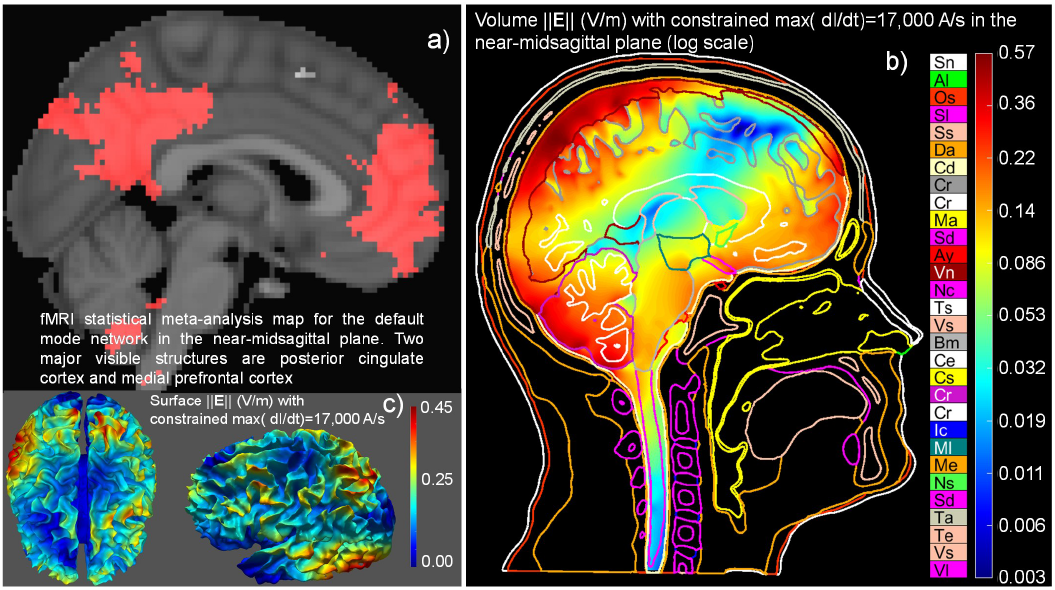
a) Meta-analysis map of DMN [70], [71] in a near-midsagittal plane. b) Beamforming attempt after approximate registration of fMRI voxel data to a Sim4Life segmentation of a subject under study. c) The corresponding electric field just outside of the white matter surface, displayed on the white matter surface.

#### Example #4. Replication of endogenous cortical activity patterns

The final example demonstrates synthesis of brain stimulation fields that emulate endogenous traveling alpha waves generated by thalamocortical oscillators [72], [73] and the Rolandic mu rhythm [74], [75], with the goal of enabling oscillation-synchronized TMS for depression treatment [44]– [46]. The ROI encompasses the entire neocortex, with the white-matter surface serving as the optimization target. We reproduced experimentally measured alpha-wave propagation in epileptic patients [76] by assuming the normal cortical electric field to be proportional to the cortical dipole density. Experimental data from [76] and one of our segmented head models were used to determine the coil weights at successive time points. Once the subject-specific basis functions are precomputed, this optimization requires less than 1 ms per time step, making prospective closed-loop EEG feedback feasible. Fig. 13 shows the resulting normal electric field just outside the white-matter surface at successive time points, normalized to an alpha frequency of 10 Hz. Consistent with the experimental observations [76], the stimulation pattern propagates from higher-order anterosuperior regions toward the occipital cortex, while activity in the somatosensory cortex propagates from associative regions toward the primary sensory cortex. Although the color scale saturates at ± 0.1 V/m, local field strengths reach approximately 0.2 V/m for a maximum individual-coil dI/dt of 0.017 A/*µ*s and 0.6 V/m for the experimentally demonstrated limit of 0.05 A/*µ*s.

**Fig. 13.**
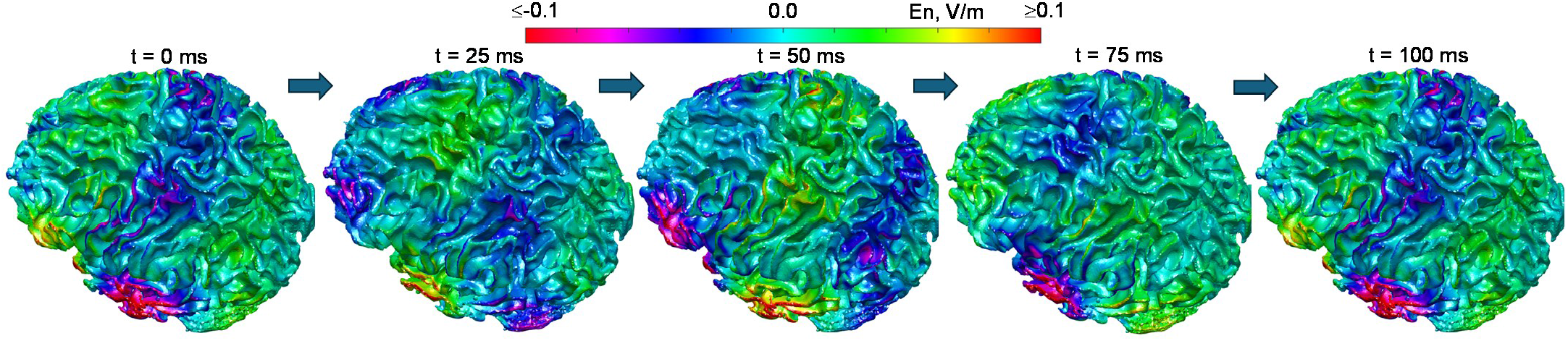
Normal component of the alpha- induced stimulation electric field just outside the white matter surface. This traveling wave was directly generated by the 102-coil array with differently evolving coil phases. The wave propagates almost exactly following temporal evolution of the density of electrical cortical dipoles extracted from the experimental data [76].

## IV. Discussion

### A. Advantages of the present multichannel whole-head array approach

The contribution of this work lies not in any single capability, but in the combination of capabilities enabled by the proposed 102-channel array. Because each channel generates an independently programmable, software-defined waveform, the array functions as a reconfigurable rather than a fixed-geometry stimulator. The same hardware can synthesize sinusoidal, multi-harmonic, or pulsed waveforms, impose arbitrary spatial and temporal phase offsets across the cortex, and stimulate multiple regions simultaneously. The proposed system might therefore represent a distinct neuromodulation modality: a low-power, whole-cortex, steerable LI-TMS array.

As set out in the Introduction, the array reaches the same cortical field regime as multichannel tACS without requiring scalp contact. It also differs from existing multichannel TMS systems, which provide electronic steering at suprathreshold intensities over cortical regions approximately 30 mm in diameter; the present array instead trades stimulation intensity for whole-cortex coverage at subthreshold field levels.

The multi-harmonic, phase-controlled waveform capability further positions the array concept for emerging temporal-interference paradigms [77], [78], which require precisely the independent per-channel frequency and phase control the system was designed to provide. Finally, because the array is non-contact, the scalp remains free for concurrent recording: a textile dry-electrode EEG cap worn beneath the helmet [79],[80] would enable near-real-time feedback and, ultimately, phase-locked closed-loop stimulation synchronized to ongoing cortical oscillations across the whole head.

### B. Limitations of the present approach

As a purely inductive, non-contact technology intended for benchtop or home use, the proposed system has several limitations and technical challenges identified during prototype development. First, unlike electrode-based stimulation, it cannot reproduce transcranial direct-current (DC) stimulation or alternating-current stimulation. In addition, the system requires considerably more sophisticated power electronics, computer hardware, and control software than conventional electrode-based tES systems. Although its operating power is substantially lower than that of conventional TMS, coil heating may still become significant at higher electric-field amplitudes, particularly during prolonged continuous-wave stimulation intended to emulate temporal interference when the current in individual coils exceeds approximately 2–3 A. Finally, the present prototype remains relatively large and heavy: the helmet weighs approximately 2.3 kg (Fig. 2), comparable to a fully equipped modern U.S. Army combat helmet [81], while the complete 102-channel driver, including the battery, weighs approximately 8 kg and occupies a volume comparable to that of a large shoebox.

Several of these limitations are being addressed through further hardware optimization. Helmet weight may be reduced by decreasing the helmet wall thickness (currently about 3 mm) and attachment method. Further improvements under development include enhanced heat dissipation and a lightweight insulating liner on the inner helmet surface to improve wearer comfort.

### C. Value of constructive and destructive interference

Both constructive and destructive interference among the individual coils are exploited during array operation. Constructive interference enables cortical electric fields substantially larger than those produced by individual coils and may facilitate deeper stimulation. In Figs. 10–13, peak cortical fields do not exceed 0.5 V/m (Fig. 11c) under the constraint max(dI/dt)≤ 0.017 A/*µ*s. The current hardware supports max(dI/dt) ≤ 0.05 A/*µ*s, although the positive pulse duration is then reduced from 2.25 ms (the original Re5 pulse) to approximately 1 ms. Under these conditions, theoretical peak cortical fields of up to 1.5 V/m are achieved despite a maximum dI/dt that is only about 0.1% of that used in suprathreshold TMS.

The explanation is that the fields generated by individual coils combine constructively. Fig. 14a illustrates left DLPFC targeting with medial-lateral field polarization, where eight small coils (four driven in phase and four in antiphase) effectively form a large figure-8 coil, producing a stronger field between its two loops. By exploiting constructive interference, the array may also generate field patterns resembling those of TMS H-coils [82] and potentially improve stimulation of deeper structures such as the hippocampus, whose potentiation has been implicated in antidepressant effects [83], [84]. Fig. 14b shows an example of hippocampal targeting with the present array, again using the constraint max(dI/dt)≤ 0.017 A/*µ*s.

**Fig. 14.**
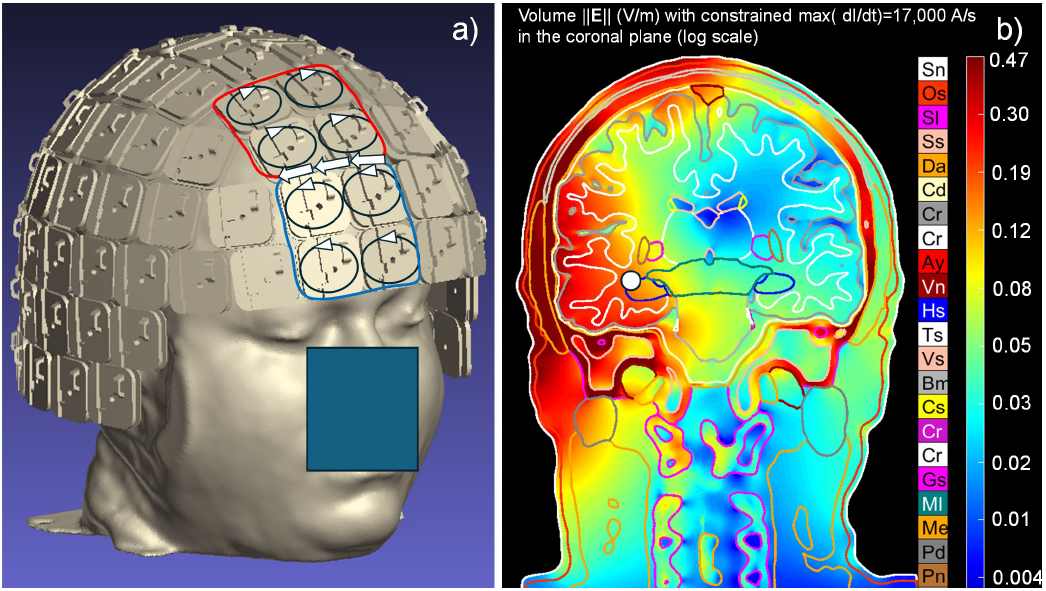
a) Example of synthesis of a large ‘shape-eight’ coil using eight individual coils. b) Beamforming for the left hippocampus (0.17 V/m at target – white sphere); with max(dI/dt)≤0.017 A/***µ***s.

### D. Applicability to temporal interference

Temporal interference (TI) stimulation delivers sinusoidal carrier waves at two slightly different high frequencies (e.g., 2 kHz and 2.01 kHz) through scalp electrodes. The intrinsic lowpass filtering of the neural membrane prevents neural activity from following the high-frequency electric fields, so that it follows only the 10 Hz envelope created by the interference of the two carriers [77]. TI has been shown to deliver envelope amplitudes of approximately 0.26 V/m to deep structures such as the hippocampus at 1 mA per electrode pair, some 44% larger than the 0.18 V/m measured in the overlying cortex, in both modeling and measurements in a human cadaver [85]. The present array can deliver an independently specified current waveform to each channel at kilohertz frequencies, which could enable TI stimulation with updated software. Because the stimulation is inductive, less current is driven through the scalp and cerebrospinal fluid than in electrode-based TI, which may permit higher stimulation amplitudes at depth.

## V. Conclusions

We have demonstrated a whole-head, low-intensity 102-channel TMS array featuring software-defined, individually controlled coils and a BEM-FMM-based beamforming framework for spatial and spatiotemporal electric-field targeting. In simulation, the array reproduced the electric and magnetic fields of the clinically validated Re5-NTS device with an accuracy of approximately 2%, enabled focal targeting of the left dorsolateral prefrontal cortex (DLPFC) with a controllable focality–intensity trade-off, engaged the default mode network, and synthesized a traveling alpha wave consistent with experimentally observed cortical propagation patterns. Direct electric-field measurements obtained with a 12-channel prefrontal subset showed good agreement with simulation, while thermal testing confirmed operation within an acceptable temperature rise range for both pulsed and continuous-wave (CW) stimulation protocols.

By combining the fields generated by multiple low-power coils, the array achieves cortical electric-field strengths comparable to those of transcranial electrical stimulation (tES) without requiring direct scalp contact and while operating at current slew rates substantially lower than those typically used in conventional TMS systems.

## Acknowledgment

The authors gratefully acknowledge valuable discussions with Drs. Aapo Nummenmaa (Massachusetts General Hospital), Søren Bøgevig (Re5 ApS), Derek Drumm (Worcester Polytechnic Institute), and Marom Bikson (City College of New York). They also gratefully acknowledge the hardware support provided by Re5 ApS (Frederiksberg, Denmark).

